# Task goals dynamically reconfigure neural working memory representations

**DOI:** 10.64898/2026.01.19.700420

**Authors:** Frida A.B. Printzlau, Olya Bulatova, Michael L Mack, Keisuke Fukuda

**Author notes:** Corresponding author: Frida Printzlau, Department of Psychology, Faculty of Arts and Sciences, University of Toronto, 100 St George St, Toronto, ON, M5S 3G3. Equal contribution (Michael L Mack, Keisuke Fukuda).

## Abstract

A hallmark of human intelligence is the ability to flexibly shift between tasks and modify our behaviour according to current goals and context. Working memory (WM) is thought to be critical for this cognitive flexibility. A key unanswered question is how neural representations of WM are configured to support different task goals. Thirty-four participants performed a shape WM task with different task rules in interleaved blocks: delayed match-to-sample (DMS) and delayed match-to-category (DMC), while electroencephalography (EEG) was collected. Our results show that stimuli from a circular shape space can be decoded from EEG across the memory delay. Neural stimulus patterns initially generalised across tasks but reconfigured to integrate task and category information and became task-dependent in the later delay. In the DMC task, neural patterns for shapes within a category became more similar than across categories, controlling for their physical distance. This category-dependent shift was associated with greater task-dependency in the neural stimulus representation. Together, our results reveal a dynamic reconfiguration of neural WM representations over the course of the delay period to support distinct task demands.

**Significance statement:** As humans, we regularly move between tasks and change our behaviour according to the situation. This cognitive flexibility critically depends on our working memory, the ability to keep information in mind in the short term. It remains poorly understood how working memory configures mental representations to support different tasks. In this study, participants performed two tasks that probed working memory for visual details or object category respectively. Neural representations were initially similar irrespective of task but became more task-dependent over time in memory. The task-dependence in the neural representation was explained by a neural category bias that emerged during the memory delay. Our results demonstrate how neural representations integrate prior knowledge with stimulus representations in a task-dependent manner, supporting flexible cognition.

---

As humans, we are very skilled at performing a range of cognitive tasks and flexibly shifting our behaviour depending on current goals and context. For example, we may go to the store to buy our favourite type of chocolate chip ice cream, but upon learning they are sold out, shift our search to the broader category of ice cream. Working memory (WM), the ability to keep information in mind over the short term, is thought to be critical for this type of flexible cognition^1–3^. Here, we ask how representations in WM are configured to flexibly support distinct task demands.

Traditionally, theories of WM have been most concerned with the retrospective mechanisms by which a detailed stimulus representation is maintained across a delay. Influential early models, inspired by observations in non-human primates, proposed stimulus-specific persistent delay activity in prefrontal cortex (PFC) as a candidate mechanism for WM function^4–7^. More recently, it has become clear that WM delay activity is highly dynamic and influenced by past experience as well as future goals^8–13^. For example, Sarma et al. (2016) showed that delay activity in posterior parietal cortex (PPC) and PFC varied depending on whether monkeys were tasked with delayed match-to-sample (DMS) or match-to-category (DMC) judgements of motion stimuli^14^. Neuroimaging studies in humans have similarly shown task differences in object representations along the ventral visual stream and PFC depending on whether participants made judgements about visual or conceptual features of the same objects^15,16^.

Sorting information into meaningful concepts is crucial for making decisions and drawing inferences. Behaviourally, category learning has been linked to changes in perceptual sensitivity along category-relevant dimensions^17^. For example, items from different categories may be perceived as more distinct and items from the same category as more similar following learning^18,19^. Similar changes have been observed in the brain. Early macaque studies showed abstract category signals emerging in PFC and PPC following learning of arbitrary category rules^20–22^. Additionally, learning can trigger neural enhancement of diagnostic category features as well as compression of neural representations to selectively code for relevant features depending on the current category rule^23–28^. Finally, recent human neuroimaging studies have found widespread category-dependent biases in stimulus representations including in early visual cortex^29–31^.

The reach of category biases extends beyond perception to similarly impact WM^32–36^. The influence of category knowledge on WM grows larger with longer delay periods^37,38^, yet it remains unknown whether biases emerge at the level of WM delay representations or during decision-making or both. This makes it important to understand at what stage of processing category priors influence WM representations and how those are affected when category rules are flexibly applied. To date, most studies investigating task-and category-dependent neural processing in humans have used fMRI, leaving the temporal dynamics of these effects unexplored^15,16,23,25,26,31,39^. This is an important gap to fill given the strong evidence for dynamic coding in WM showing rapid changes to the neural patterns supporting WM between encoding and delay periods, within a few hundred milliseconds^9–11,40^.

In this EEG study, we asked whether WM delay representations for the same stimuli were modulated by current task demands. Participants first learned to sort unfamiliar shapes from the validated circular shape (VCS) space^41^ into arbitrary categories, minimising interference from pre-existing category knowledge. Next, they performed a WM task, with DMS or DMC judgements in different, interleaved task blocks. We recorded EEG during the WM task to track the dynamics of stimulus-, category and task-level representations over the course of the memory delay. If current task goals reconfigure WM representations to support prospective behaviour, the neural stimulus pattern may differ as a function of task^16^.

To preview our results, although stimulus decoding quality was similar across the two tasks, the neural pattern underlying these stimulus representations became task-dependent in the late memory delay. Representational similarity analyses (RSA)^42^ reveal that neural signals dynamically tracked stimulus-, category-and task-level information over the course of the delay, with stronger category signals in the DMC task. Finally, this was accompanied by category-dependent shifts in the stimulus representation in the DMC task, where the neural representations for same-category stimuli became more similar than for different-category stimuli, predicting the task-dependency in stimulus representations. Together, our results suggest that recent learning history can reconfigure neural WM representations to support current task goals. Specifically, WM flexibly integrates prior knowledge with stimulus representations in a goal-dependent manner in preparation for a forthcoming decision.

## 1. Results

### 1.1 Behaviour

#### 1.1.1 Category learning

Participants first completed a category learning task (Figure 1A; see *Materials and Methods*) asking them to categorise twelve shapes from the VCS space^41^ according to an arbitrary category boundary (counterbalanced across participants). Participants reported whether the shape was a ‘Fuboo’ or a ‘Juboo’ using the keyboard and received feedback after each response. Participants completed eight learning blocks of 24 trials.

**Figure 1.**
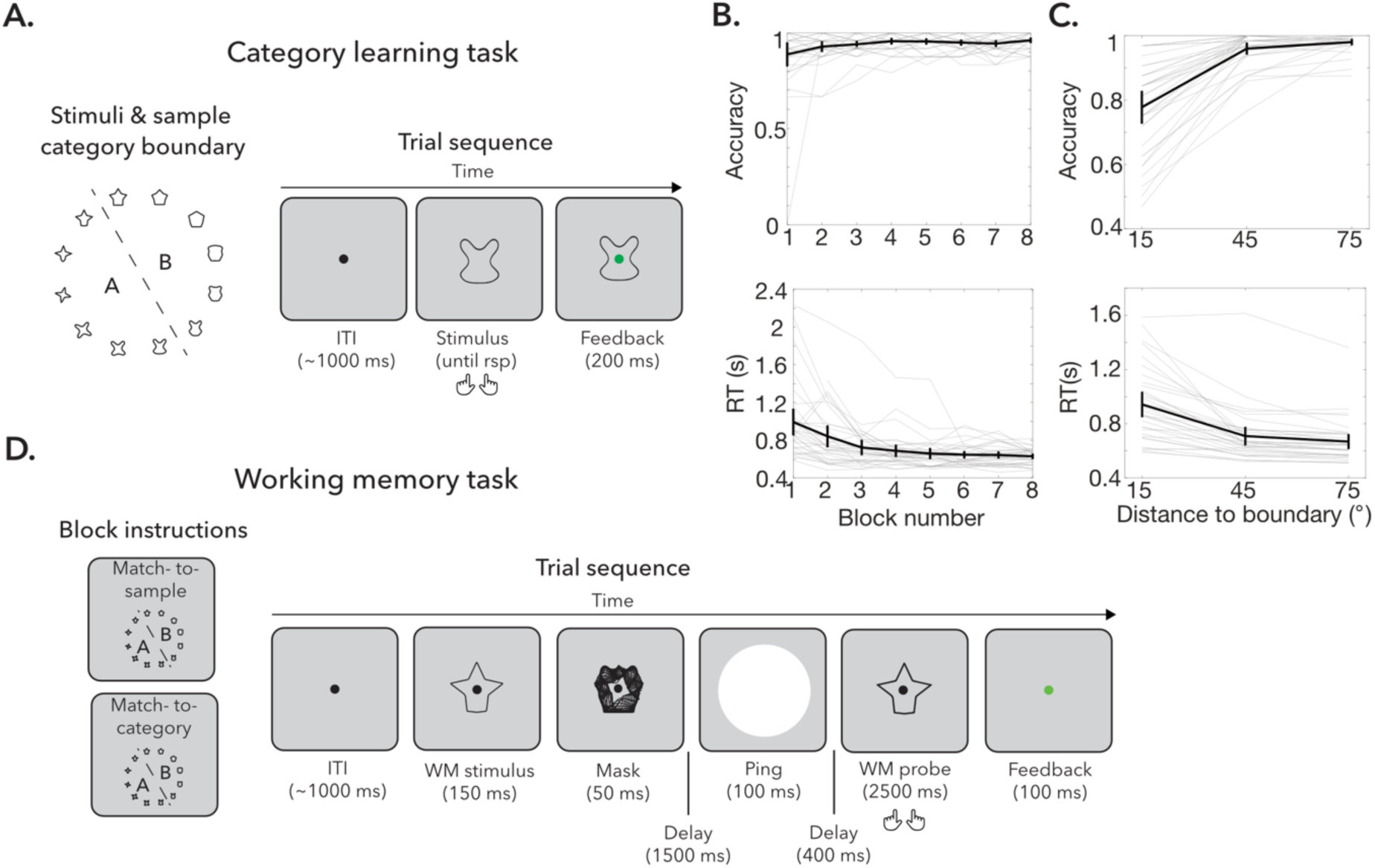
Task design. **A**. Stimuli and trial sequence of category learning task. Participants viewed the 12 shape stimuli and category boundary prior to starting the learning task (left panel). On each trial, participants were presented with a shape and asked to respond which category it was a member of. They received feedback after their response. **B.** Category learning accuracy (top) and RT (bottom) as a function of block. Participants completed 8 blocks of learning. Participants were required to achieve 90% accuracy on at least one learning block to proceed to the working memory task. Error bars show 95% confidence intervals (CI). Grey lines show individual participants. **C.** Category learning accuracy (top) and RT (bottom) as a function of distance to the category boundary. **D.** Working memory task trial sequence. Prior to each trial block, participants received instructions whether the next block would be a match-to-sample task or match-to-category task. They were also reminded of the category boundary at the start of each block. On each trial, they were presented with a WM stimulus (of 36 total) followed by a mask. After a 1500 ms delay a ping stimulus appeared followed by another brief delay before the probe was presented. Participants were asked to compare the probe to the memory target according to the currently relevant task rule and submit their response via a button press. They received feedback after each response. Stimuli not drawn to scale.

All participants scored above 90% accuracy in at least one learning block, meeting the inclusion threshold (Figure 1B). Learning performance improved during training, with a main effect of learning block on accuracy (F_7,231_=3.69, *p_GG_corr_*=.033, 17_p_^2^=.101) and RT (F_7,224_=18.90, *p_GG_corr_* <.001, 17_p_^2^=.371).

Learning is thought to be more difficult near a category boundary. We found a main effect of the angular distance to the category boundary on both accuracy (F_2,66_=71.17, *p_GG_corr_*<.001, 17_p_^2^=.683) and RT (F_2,66_=53.28, *p_GG_corr_* <.001, 17_p_^2^=.618; Figure 1C), with lower accuracy and higher RTs for shapes near the boundary relative to shapes closer to the centre of the category (Accuracy: 15° vs 45°: t_33_=-8.97, *p*<.001, *d*=-1.61, 95% CI=-.22 to-.14; 45° vs 75°: t_33_=-2.63, *p*=.013, *d*=-.452, 95% CI=-.036 to-.005; RT: 15° vs 45°: t_33_=6.87, *p*<.001, *d*=.960, 95% CI=.164 to.302; 45° vs 75°: t_33_=3.98, *p*<.001, *d*=.219, 95% CI=.020 to.061).

#### 1.1.2 Working memory

Following learning, participants completed a WM task (Figure 1D; see *Materials and Methods*). On each trial, participants memorised a shape from the VCS space (36 possible; chosen at random without replacement within each block) for a subsequent memory judgement. After a two-second delay, a probe appeared and participants compared the probe to the memory target according to one of two task rules, on different task blocks: match-to-sample (DMS) or match-to-category (DMC). In DMS blocks participants responded ‘match’ if the probe was an exact match to the memory target (50% of trials) and ‘non-match’ if the probe was any other shape (50%). In DMC blocks, participants responded ‘match’ if the probe belonged to the same category as the memory target (50% of trials) and ‘non-match’ if the probe belonged to the opposite category (50%).

Participants were instructed of the task rule and shown the category boundary before each block. There were 864 trials in total, divided into 24 blocks of 36 trials.

WM performance was generally better on DMS than DMC trials (Figure 2A-B), reflected in higher accuracy (t_33_=4.02, *p*<.001, *d*=.698, 95% CI=.015 to.047) and faster reaction times (RT; t_33_=-13.10, *p*<.001, *d*=-1.65, 95% CI=-.200 to-.145).

**Figure 2.**
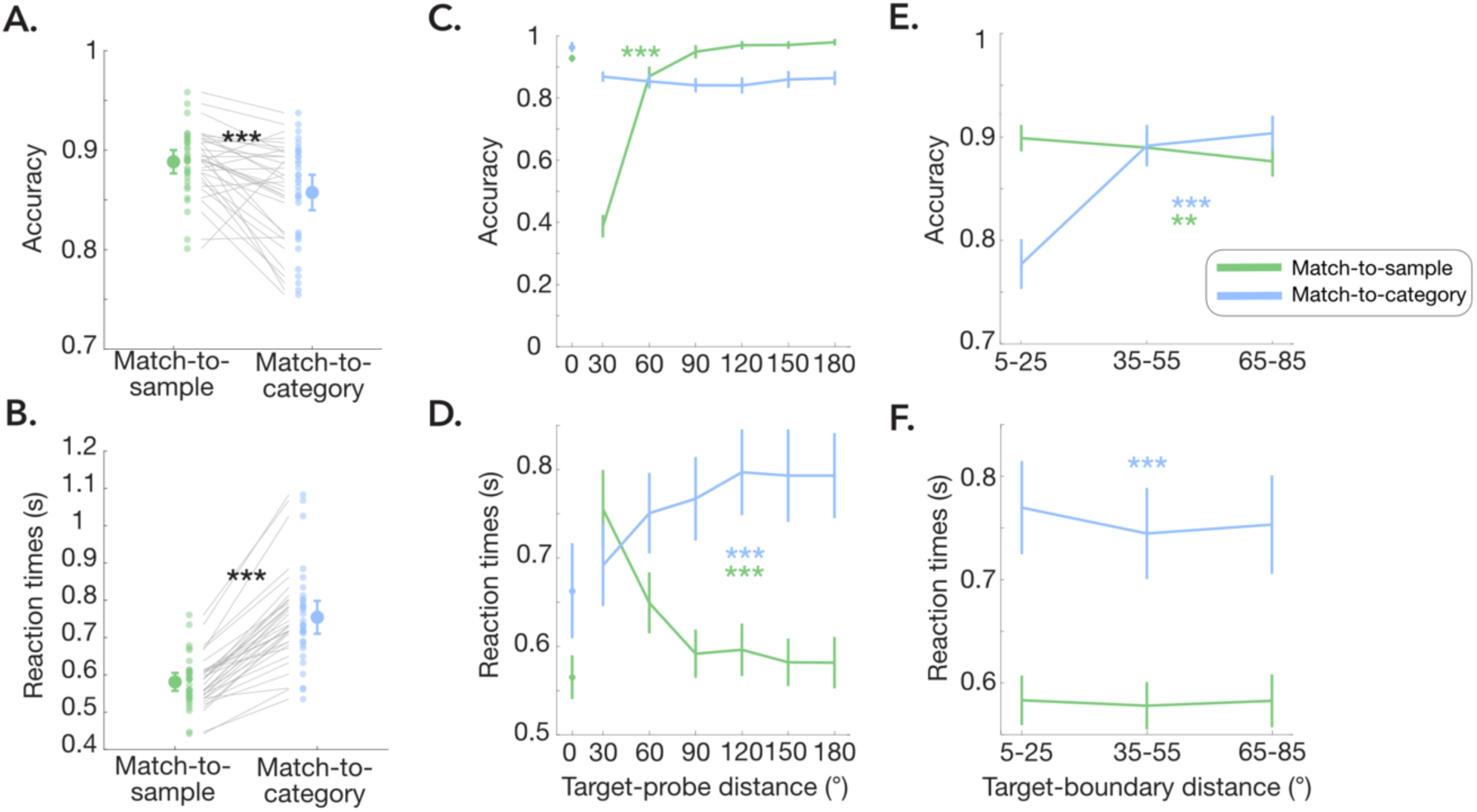
Behavioural performance on the WM task. **A**. WM accuracy on DMS task (green) and DMC task (blue). Large circle represents mean and error bars show 95% confidence intervals (CI). Small dots show individual participants and lines connect individual participants. **B.** Same as A but for reaction times**. C.** Accuracy as a function of the angular distance between the target and the probe. Error bars show 95% CI **D.** Same as C but for reaction times**. E.** Accuracy as a function of the angular distance between the target and the category boundary**. F.** Same as E, but for reaction times.

We examined performance as a function of the angular distance between the target and the probe, relevant to DMS judgements (Figure 2C-D). On DMS trials, accuracy increased (F_5,165_=644.9, *p_GG_corr_*<.001, 17_p_^2^=.951) and RTs decreased (F_5,165_=63.02, *p_GG_corr_*<.001, 17_p_^2^=.656) with greater target-probe distances (excluding target-probe distance of zero). On DMC trials, we did not find a main effect of target-probe distance on accuracy (F_5,165_=2.13, *p*=.065, 17_p_^2^=.061), but RTs increased with greater target-probe distances (F_5,165_=27.49, *p*<.001, 17_p_^2^=.455), suggesting that participants used target-probe similarity in their category decisions, to some extent.

Next, we looked at performance as a function of the angular distance of the target from the category boundary, relevant to DMC judgements (Figure 2E-F). On the DMC task, we found a main effect of target-boundary distance on both accuracy (F_2,66_=125.57, *p_GG_corr_*<.001, 17_p_^2^=.792) and RTs (F_2,66_=7.86, *p*<.001, 17_p_^2^=.192) as expected, with lower accuracy and higher RTs near the category boundary relative to category centre. On the DMS task, there was a main effect of target-boundary distance on accuracy (F_2,66_=6.51, *p*=.003, 17_p_^2^=.165), but not on RT (*p*=.356), with higher accuracy near the category boundary relative to the centre. This was surprising because target-boundary distance was irrelevant to DMS decisions and suggests that category structure nevertheless informed DMS judgements.

A follow-up analysis showed that the effect of target-boundary distance on DMS accuracy was driven by the difficult non-match trials, when the distance between the target and the probe was small (0<probe<90°; F_2,66_=11.35, *p*<.001, 17_p_^2^=.256), rather than easy non-match trials or match trials (p>.6; See *Supplementary material*; Figure S1A). This suggests items near the category centre may be encoded with lower precision than items near the boundary or be systematically biased. To further test if DMS judgements were biased by category structure, we checked if false alarms were more likely for non-match probes that were toward (attractive bias) or away (repulsive bias) from the category centre (Figure S1B). Limiting our analysis to trials where the target and probe were within the same quadrangle, we found higher accuracy on ‘toward’ than ‘away’ trials (t_33_=3.07, *p*=.004, *d*=.551, 95% CI=.025 to.124), meaning people false alarmed more to non-targets that were away from the category, consistent with a repulsion away from the category centre.

### 1.2 EEG

#### 1.2.1 Decoding stimulus shape

We collected EEG during the WM task using a 32 channel BrainVision system (27 EEG channels used for analyses)^43^. If the activity measured in the EEG channels contained information about the target shape in memory, we would expect the neural pattern similarity (measured using inverse Mahalanobis distance) to be higher for shapes that were close together in the circular shape space and lower for shapes that were far apart (see *Materials and Methods*)^44^. We limited our stimulus decoding analyses to correct trials to ensure the target was encoded well (subsampling ensured equal trial numbers across conditions). A follow-up analysis showed no difference in decoding quality between correct and incorrect trials (see *Supplementary Material*; Figure S2).

Figure 3A shows the neural pattern similarity over time after stimulus onset for shapes sorted relative to the memory target. Neural similarity following memory encoding was higher for near shapes and lower for shapes that were distant in the circular shape space. Summarising pattern similarity into a measure of decoding quality by taking the cosine weighted mean^44,45^, we find that decoding quality was significantly above chance (zero) from stimulus presentation until shortly before probe presentation (0.03-1.85 s; cluster-corrected *p*<.001; Figure 3B).

**Figure 3.**
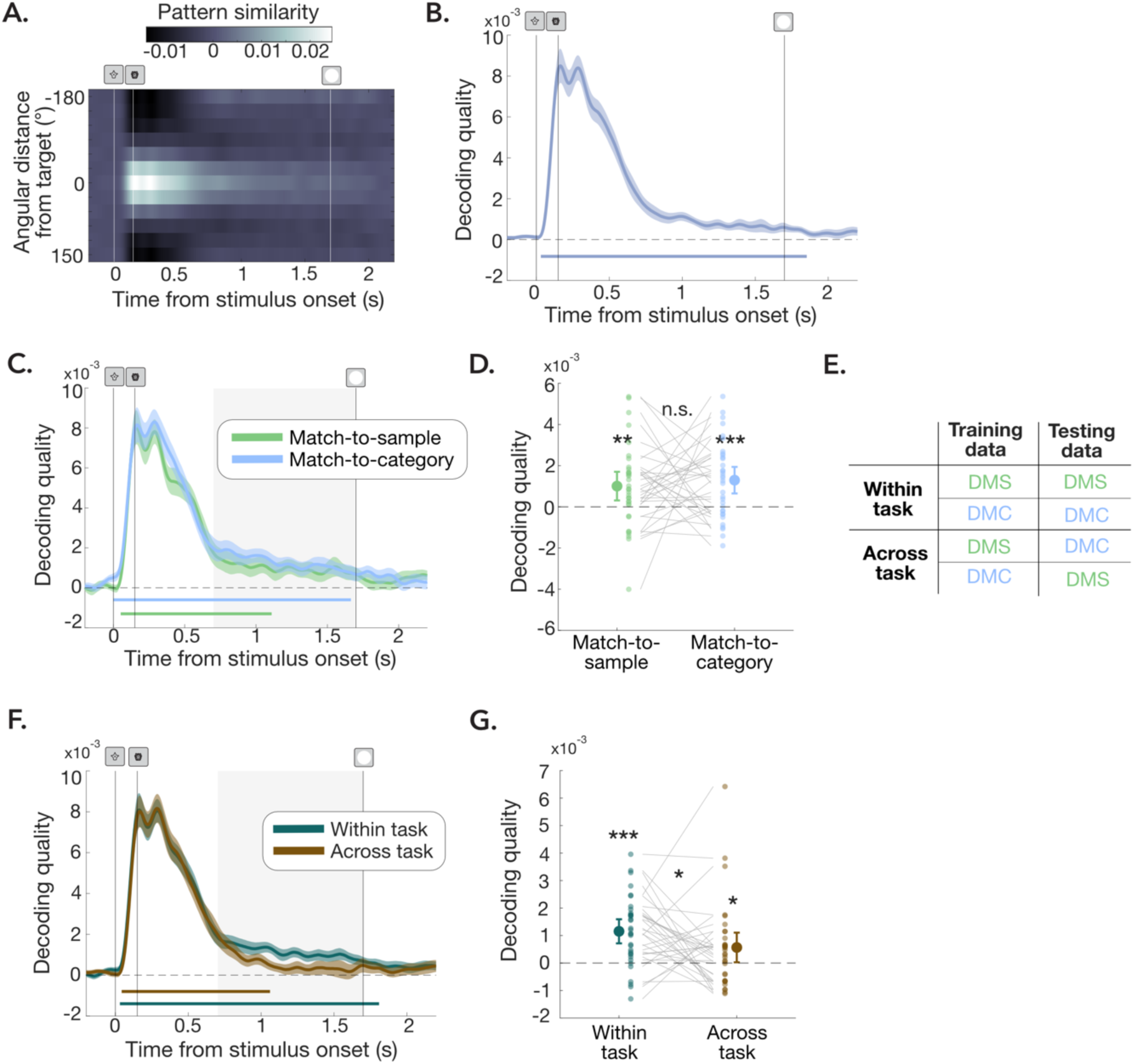
Decoding shape in WM. **A.** Neural pattern similarity (negative cross-validated Mahalanobis distance) for shapes relative to target over time after stimulus presentation. **B.** Decoding quality over time after stimulus onset. Error bars show standard error of mean (SEM). Coloured bars show significant time points after cluster correction (p<.001). **C.** Same as B, but separately for the DMS task (green) and DMC task (blue). Grey box shows late delay window shown in D. **D.** Mean decoding quality in late time window (0.7-1.7 s after stimulus presentation). Large dots show group mean. Error bars show 95% CI. Small dots show individual participants. Grey lines connect individual participants. **E.** The cross-task decoding procedure that either involved training and testing on data from the same task (within task decoding) or training on data from one task and testing on data from the other task (across task decoding). **F.** Within task (cyan) and across task (brown) decoding quality over time from stimulus onset. Otherwise, same as B. **G.** Mean decoding quality in late time window (0.7-1.7 s) separately for within-task (cyan) and across-task (brown) conditions. Otherwise, same as D. \**p*<.05, \*\**p*<.01, \*\*\**p*<.001.

A strong visual impulse or ‘ping’ has previously been shown to improve the decodability of orientation stimuli from EEG^44,46^. We presented a ping late in the memory delay (1.7 s) but did not find any boost in decoding quality following ping onset (Figure 3B). For that reason, we focus our analyses on the ‘clean’ delay period before ping onset.

Initial decoding quality was high and decreased to a lower level in the later delay (Figure 3B). We defined an early and a late time-window of interest based on the shape of the overall decoding time-course. We computed the derivative of mean decoding quality over time across participants and set a threshold of +/– 0.01 of the derivatives to identify the time-point at which decoding accuracy plateaued (time=0.7 s). Consequently, we focus our statistical analyses on an early time-window from 0 to 0.7 s and a late time-window from 0.7 to 1.7 s.

#### 1.2.2 Stimulus decoding quality is similar for both tasks

We repeated the decoding analysis specified above with minor adjustments (see *Materials and Methods*) separately within each task (Figure 3C). In both tasks, stimulus decoding was significantly above chance from shortly after stimulus onset for at least 1 s (DMS: 0.05 to 1.11 s; DMC: 0 to 1.67 s; cluster-corrected *p*<.001). Stimulus decoding was significant for both tasks in both the early (DMS: t_33_=11.60, *p*<.001, *d*=1.94, 95% CI>.004; DMC: t_33_=11.85, *p*<.001, *d*=1.99, 95% CI>.005) and late time windows (DMS: t_33_=2.94, *p*<.001, *d*=.493, 95% CI>.428*10^-3^; DMC: t_33_=4.09, *p*<.001, *d*=.696, 95% CI>.760*10^-3^). Interestingly, we find no difference in decoding quality between the two tasks in either time window (*p>*.358; Figure 3D). A searchlight analysis also revealed a similar distribution of channels contributing to stimulus coding across the two tasks (*Supplementary Material*; Figure S3).

#### 1.2.3 Task-dependent stimulus code

While stimulus decoding quality was similar across tasks, it is possible that the neural format underlying these shape representations differs depending on task^16^. If so, we would expect decoding quality to be better when training and testing the decoder within the same task (within-task decoding) compared to training the decoder on one task and testing on the other (across-task decoding; Figure 3E). On the other hand, if the neural pattern coding for shape is shared across tasks, we would not expect any cost of cross-decoding.

Because we found no task-differences in stimulus decoding quality, we averaged DMS and DMC within-and across-task decoding for improved power (see *Supplementary Material* Figure S4A for cross-decoding results separately by task).

Both within-and across-task decoding quality was significant in the early time-window (Within: t_33_=13.20, *p*<.001, *d*=2.21, 95% CI>.0044; 0.03 to 1.81 s; cluster-corrected *p*<.001; Across: t_33_=11.13, *p*<.001, *d*=1.86, 95% CI>.0042; 0.04 to 1.06; cluster-corrected *p*<.001) and there was no difference between conditions (*p*>.391). When directly comparing within-and across-task decoding quality in the late time-window (0.7-1.7 s; Figure 3F), both were above chance (Within: t_33_= 5.36, *p*<.001, *d*=.897, 95% CI>.789*10^-3^; Across: t_33_= 2.14, *p*=.020, *d*=.358, 95% CI>.118*10^-3^), but within-task decoding quality was significantly greater than across-task decoding quality (t_33_= 1.85, *p*=.036, *d*=.408, 95% CI>.510*10^-4^).

This cost of cross-decoding indicates that the neural pattern underlying the shape representation is task-dependent.

A searchlight analysis (*Supplementary material*; Figure S4B) indicated greater contribution of anterior channels to within-task decoding quality in the late delay compared to across-task decoding that was specific to posterior channels. However, only one anterior channel (Fz) showed greater within>across task decoding after correcting for multiple comparisons (*p*<.05/27).

#### 1.2.4 Tracking neural geometry of stimulus-, category-, and task-level information

We used RSA to track the neural geometry coding for stimulus-, category-and task-level information over time^42^. To compute representational dissimilarity matrices (RDM), we first rotated the shape space to align the category boundary across participants. The 36 shapes were sorted into 12 shape bins for each task, creating a total of 24 conditions. For each time-point, we calculated the mean EEG activity for trials belonging to each condition and computed the pairwise neural dissimilarity (Mahalanobis distance) between each of the conditions (see *Materials and Methods*). Given that stimulus decoding quality was similar for correct and incorrect trials (*Supplementary material*; Figure S1), we included all trials in the RSA analyses to improve power. We plotted the mean RDMs for each time-window of interest (Figure 4A). Figure 4B illustrates what a hypothetical stimulus code, category code, and task code should look like. To better illustrate the representational geometry, we also plotted the distances between conditions in two and three dimensions, using multidimensional scaling (MDS; Figure 4C & D).

**Figure 4.**
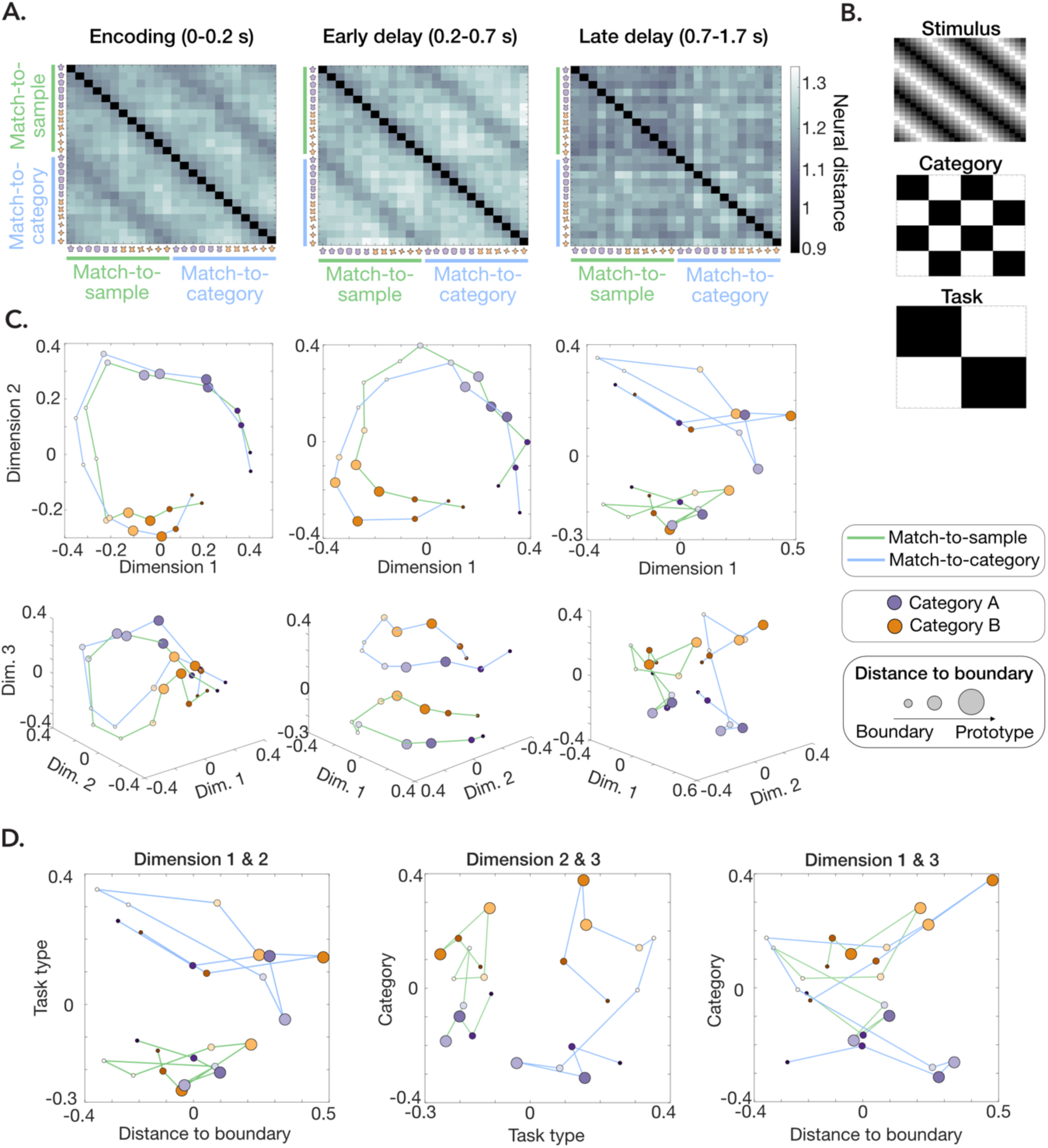
Neural geometry of stimulus-, category-, and task-level information**. A.** Neural RDMs during encoding (left; 0-0.2 s after stimulus presentation) the early delay period (middle; 0.2-0.7 s) and late delay period (right; 0.7-1.7 s). Each cell in a matrix corresponds to the Mahalanobis distance between a 30 deg bin of stimulus values and all other shape bins sorted according to task (DMS in green and DMC in blue) and category (purple vs orange). **B.** Hypothetical stimulus-(top), category-(middle) and task-(bottom) structure. **C.** MDS plots illustrating the neural distances between conditions in two (top) and three (bottom) dimensions for each of the time-windows of interest. The hue of each circle represents the individual stimulus bin, the colour of the circle (purple vs orange) shows the category membership, the size of the circle shows the distance to the category boundary (small=near, large=far). Finally, the colour of the lines connecting the circles shows the task condition: DMS (green) or DMC (blue). **D.** MDS plots of the late delay period (07-1.7 s) plotting the first and second dimension (left), second and third dimension (middle) and first and third dimension (right), coding for distance to boundary, task type, and category membership respectively.

From visual inspection, the neural representational similarity structure matches the circular structure of the stimulus space initially after encoding (0-0.2 s) and is relatively invariant to task condition (Figure A&C, left panel). In the early delay after stimulus and mask offset (0.2-0.7), the circular structure is well-maintained for both tasks, but separates by task along a third dimension (Figure A&C, middle panel). In the late delay (0.7-1.7 s), stimulus representations cluster by task (Figure 4D; left and middle panels) and the circular stimulus structure becomes less pronounced. Instead, we start to see strong coding for the distance to category boundary (Figure 4D, left and right panels), as well as greater separation along the category-relevant axis in the DMC relative to DMS task (Figure 4D, middle and right panels).

#### 1.2.5. Greater coding for category-level information in DMC task

To formally test the extent to which neural activity coded for stimulus-and category-level information in each task, we fit a generalized linear model to each time-point of the RDM, with a graded stimulus model, a category model, and a distance-to-boundary model as predictor variables (Figure 5A; see *Materials and Methods*). The models were fit separately for each task. We extracted beta coefficients for each predictor at each time point and these were tested for deviations from zero.

**Figure 5.**
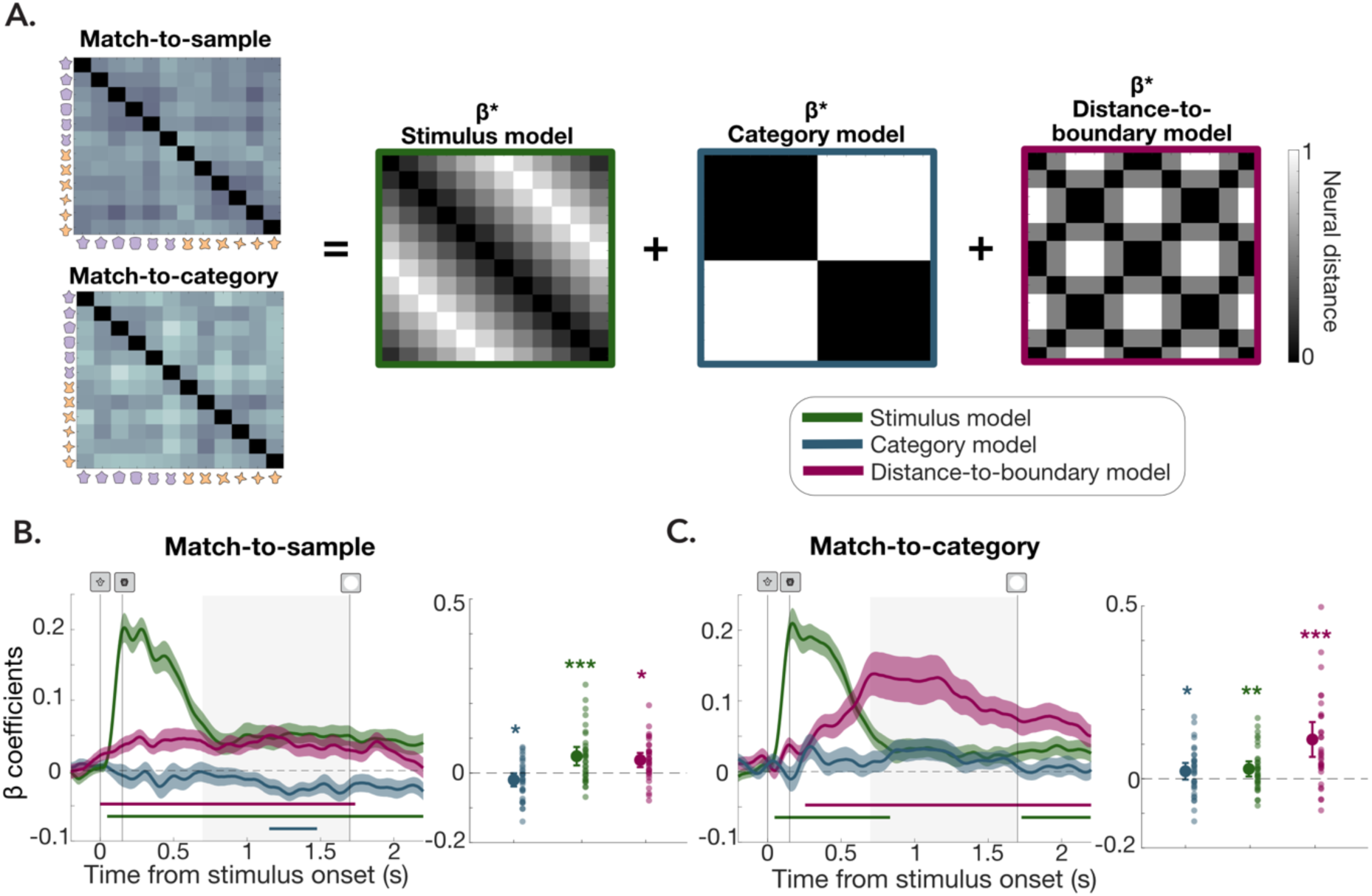
Representational similarity analysis. **A.** Illustration of stimulus model (green), category model (blue), and distance-to-boundary model (pink) that were fit to DMS and DMC neural distance data separately to obtain betas for each regressor. **B.** Regression coefficients (betas) over time for DMS task. Shaded areas show SEM. Coloured bars show significant timepoints after cluster correction (p<.05). Grey box shows time-window for averaging over late delay period (0.7-1.7 s) shown in left panel and used for statistical testing. Large dot shows mean. Error bars show 95% CI. Small dots show individual participants. **C.** Same as B, but for DMC task. \**p*<.05, \*\**p*<.01, \*\*\**p*<.001.

In both tasks, we found that beta coefficients for the stimulus coding model were significantly above zero following stimulus presentation and into the delay period, mirroring the decoding results presented above (Figure 5B&C). In the DMS task, stimulus coding remained significant for the entire delay (0.05 to 2.2 s; cluster-corrected *p*<.001). Although stimulus coding dropped in the later delay in the DMC task (0.04 to 0.83; cluster-corrected *p*<.001; 1.72 to 2.19 s; cluster-corrected *p*=.038) it remained significant in the late time-window for both tasks (DMS: t_33_= 3.71, *p*<.001, *d*=.621, 95% CI>.026; DMC: t_33_= 2.65, *p*=.006, *d*=.445, 95% CI>.010) and there was no significant difference between tasks (*p*=.217).

In the DMC task, we found significant category coding in the late delay period (Figure 5C; t_33_=1.83, *p*=.038, *d*=.306, 95% CI>.002). Category coding was significantly greater in the DMC than DMS task (t_33_=-2.83, *p*=.008, *d*=-.651, 95% CI=-.071 to-.012). In the DMS task, there was a surprising negative relationship with the category model in the late delay (1.15 to 1.48 s, cluster-corrected *p*<.042; t_33_=-2.12, *p*=.042, *d*=-.355, 95% CI=-.0386 to-.0008). This suggests that the neural pattern in the late delay was more similar for shapes within a category than across categories in the DMC task, but vice versa in the DMS task.

Interestingly, neural activity coded for the distance to the category boundary in both tasks early after stimulus onset and across the delay (DMS: 0 to 1.74 s; DMC: 0.26 to 2.2 s; cluster-corrected *p*<.001), even though the category boundary was only relevant in the DMC task. In the late delay period, distance-to-boundary coding was significantly greater for DMC than DMS task (t_33_=-3.12, *p*=.004, *d*=-.678, 95% CI=-.126 to-.027), though significant for both (DMS: t_33_=3.72, *p*<.001, *d*=.623, 95% CI>.020; DMC: t_33_=4.60, *p*<.001, *d*=.770, 95% CI>.072). This suggests that neural activity codes for category-level information in both tasks, but more so in the DMC than DMS task in the late delay.

#### 1.2.6. Category-dependent representational shift

The category model in our RSA analysis was only designed to detect a very coarse category code, corresponding to a semantic label where all shapes within a category are represented similarly. Because stimulus coding, category coding and distance-to-boundary coding were all significant in the late delay in the DMC task, we speculated that more subtle representational changes may occur as a function of category membership. For example, if the neural representation of the shape space is stretched along the axis in feature space that separates the two categories (Figure 6A)^26,28^, we would expect neural patterns to be more similar for shapes belonging to the same category than shapes from different categories, controlling for the physical distance in the shape space. We subtracted the neural distance between shapes from different categories from the neural distance between shapes from the same category, matched for the angular distance between shapes in the circular space, to get a measure of ‘category-dependent representational shift’ (Figure 6A). If the neural representation of the shape space is stretched along the category axis, we would expect this measure to be positive.

**Figure 6.**
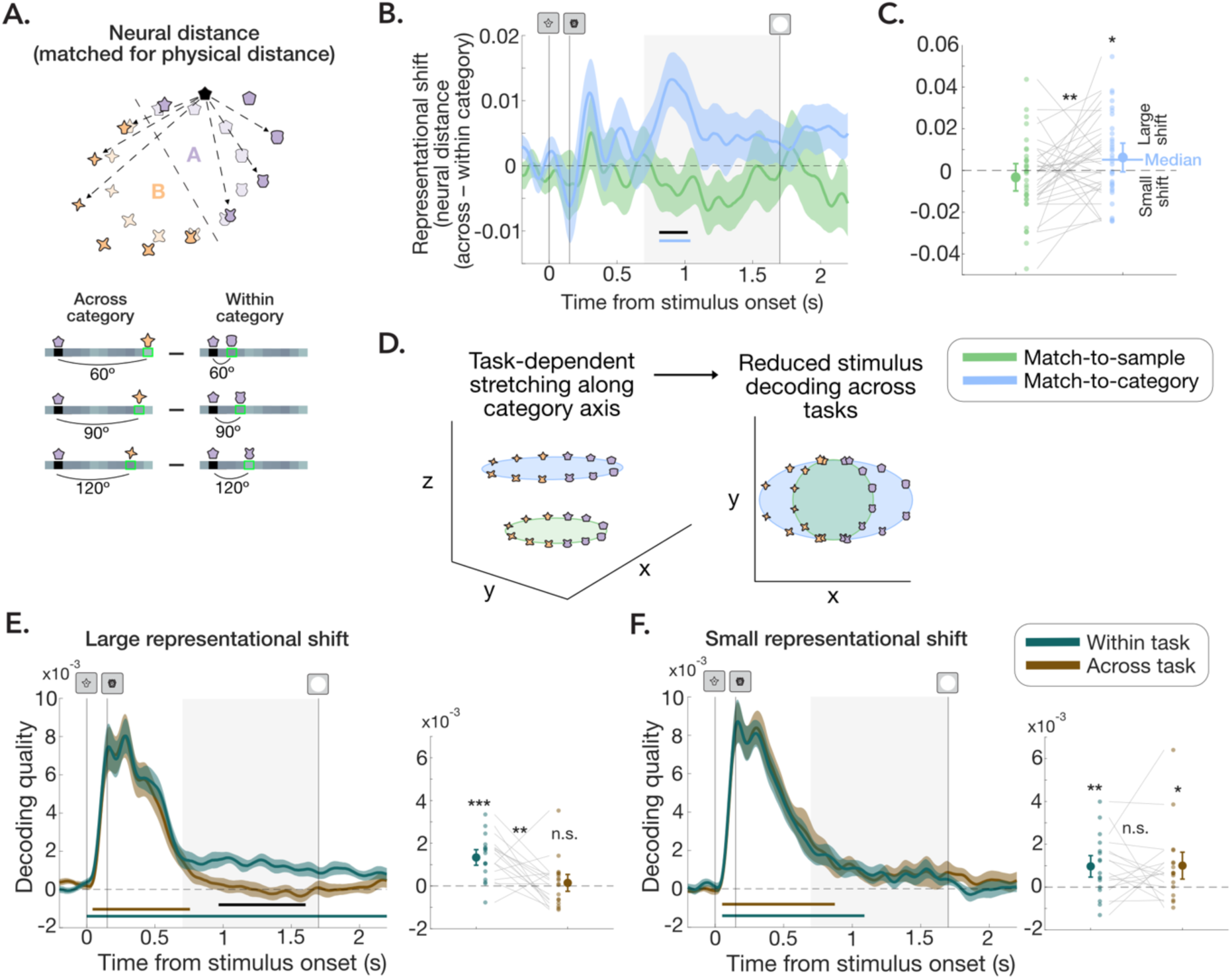
Category-dependent representational shift. **A.** For each shape, we obtained the neural distance to all other shape belonging to the same category and shapes belonging to the other category and matched these for physical distance in the circular shape space. We subtracted within-category distances from across-category distances and averaged across all distances to obtain a measure of category-dependent shift. Positive values suggest smaller neural distances within than across categories. **B.** Category-dependent representational shift over time from stimulus onset for DMS task (green) and DMC task (blue). Shaded area shows SEM. Coloured bars show significant (or near significant) timepoints after cluster correction (p<.1). Grey box shows late-delay time-window (0.7-1.7 s) plotted in C and used for statistical testing. **C.** Mean category-dependent shift in late delay period (0.7-1.7 s). Large dots show mean. Error bars show 95% CI. Small dots show individual participants connected by grey lines. Blue line shows DMC median used for median split of participants in F & G. **D.** Hypothetical MDS plots illustrating task-dependent stretching of the stimulus space along the category axis in the DMC task in three dimensions (left) and two dimensions (right). Stretching might reduce cross-decoding of stimulus across tasks. **E**. Within-task (cyan) and across-task (brown) decoding quality over time (left) and averaged over late time window (right) for participants showing category-dependent representational shift>median in DMC task. **F.** Same as D, but for participants showing category-dependent representational shift<median. \**p*<.05, \*\**p*<.01, \*\*\**p*<.001.

Neural patterns were more similar within than across categories in the late delay period in the DMC task (t_33_=1.84, *p*=.038, *d*=.308, 95% CI>.494*10^-3^; 0.81 to 1.04 s cluster-corrected *p*=.068; Figure 6B), but not in the DMS task (*p*=.844) and this representational shift was significantly greater in the DMC than DMS task in the late delay (t_33_=-2.20, *p*=.017, *d*=-.483, 95% CI<-.002; 0.81 to 1.02 s; cluster-corrected *p*=.046; Figure 6C). This suggests that there were subtle, but robust, category-dependent shifts of stimulus representations in the DMC task, as expected if the neural stimulus space is stretched along the category axis.

#### 1.2.7. Category-dependent shift predicts task dependency

Finally, we hypothesized that the category-dependent representational shift observed in the DMC task could account for the task-dependency in the neural stimulus pattern that we also observed. If the neural stimulus space is stretched along the category axis in the DMC task this should result in reduced cross-generalization of the stimulus pattern between the two tasks (Figure 6D). Therefore, we expected that participants with greater category-dependent representational shift would show greater task-dependency in their stimulus representations.

We performed a median split of participants by their ‘representational shift’ score (across–within category neural distance) in the late delay period of the DMC task (0.7-1.7 s; Figure 6C) and computed the cross-decoding results separately for participants showing large and small category-dependent representational shift (Figure 6E & F). There was a significant cross-decoding cost in the late delay period for participants showing large representational shift (t_16_=2.97, *p*=.005, *d*=1.04, 95% CI>.491*10^-^3; 0.97 to 1.61 s; cluster-corrected *p*=.011), but not for participants showing small representational shift (*p*=.530). The cross-decoding cost was significantly greater for participants showing large compared to small representational shift (t_32_=2.03, *p*=.025, *d*=.681, 95% CI>.205*10^-3^). This suggests that category-dependent shifts in the stimulus representation may contribute to the task-dependent coding of shapes we observed in the late delay.

## 2. Discussion

WM is a highly flexible system that allows us to store information for different behavioural goals. In this study we asked how WM delay representations are configured for distinct task demands. We used EEG to track the dynamics of stimulus-, category-and task-level information over time while participants performed delayed match-to-sample (DMS) or delayed match-to-category (DMC) judgements.

First, we demonstrate that the validated circular shape space^41^ can be decoded from EEG, which to our knowledge has not previously been shown. This finding motivates a fruitful avenue for researchers looking to study the neural basis of WM using unfamiliar stimuli while maintaining tight control of stimulus parameters. Second, we show that neural stimulus patterns initially generalized across tasks but became task-dependent in the later delay, evidenced from reduced stimulus decoding across tasks compared to within tasks. RSA revealed that neural activity initially coded for stimulus identity but reconfigured to integrate task-and category-level information, particularly in the DMC task. Importantly, in the DMC task, neural patterns coding for shapes within a category became more similar than for shapes across the category boundary when matched for physical distance. The resultant category-dependent stretching of the stimulus space in the late delay helps explain the task-dependency in the neural stimulus patterns. Together, our findings reveal that task demands flexibly reconfigure neural WM representations to support distinct goals.

Previous fMRI studies have shown widespread task-dependent object representations in the ventral visual stream, hippocampus and LPFC^15,16,27,31^. Our results expand on the previous literature in two important ways. First, by using EEG we were able to capture the temporal dynamics of task-dependent coding in WM showing reconfiguration of the neural patterns supporting WM within a few hundred milliseconds. Numerous studies have shown rapid changes to the neural patterns supporting WM between encoding and delay periods^9–11,40^. We elaborate on these findings by showing that these dynamics may serve to configure WM representations for the expected task. This is consistent with a dynamic coding framework of WM proposing that the neural representation in the delay phase is optimised for decision-making by changing how the network responds to new input under different task conditions^9,11^, perhaps in the form of a matched filter where new sensory input is filtered through a neural pattern coding for the memory item (DMS) or its category (DMC) to compute the relevant decision signal depending on the context^47–51^. Although the spatial resolution of EEG is limited, a searchlight analysis did show greater contribution of frontal channels to within>across task decoding quality, consistent with previous reports of stronger task-dependency at later stages of the visual hierarchy^16,39^. If the mnemonic template is configured to support decision-making through top-down signals, this might explain why we see a task-dependent reconfiguration of the neural pattern coding for the stimulus toward the end of the memory delay.

Second, by employing a parametric stimulus space we were able to track the format of the neural stimulus representations as a function of task, shedding light on the mechanisms underlying the observed task-dependency. One possibility is that task-dependent stimulus representations emerge because participants simply ‘drop’ visual details in the DMC task and instead maintain a fully abstracted code, such as a category label^15^. However, we have reasons to believe that was not the case in our study. First, overall decoding quality was similar across the DMS and DMC tasks. Our decoding metric tests if the neural activity pattern mimics the circular similarity structure of the stimulus space^44^, so we would expect decoding quality to be lower in the DMC than in the DMS task if visual details were dropped. Second, behavioural RT results in the DMC task suggests participants remained sensitive to the angular distance between the memory target and the probe, which would not be the case if they had only maintained a semantic category label.

Instead, our results suggest that the task-dependency emerged due to category-dependent shift in the neural population response in the DMC task, where neural patterns for within-category items became more similar than items from different categories, controlling for physical distance in the feature space. Our results are consistent with a previous study showing categorical biases in population orientation tuning reconstructed from BOLD and EEG. In that study, biases were found as early as V1 and within 250 ms of stimulus onset, which is earlier than what we report. However, in a follow-up DMC experiment where participants could not prepare a response until probe onset (as in our study), category biases emerged gradually over the delay period, which is more in line with our results.

Multiple mechanisms could explain the category-dependent shift in the neural population response. Models of category learning propose that category-dependent representations may emerge through selective attention to relevant feature dimensions^17,52^. This is supported by studies showing neural tuning to category-relevant feature dimensions following learning, such as stretching neural representations along category relevant stimulus dimensions and/or compression along category-irrelevant dimensions^23,25–28^. These studies employed stimuli with separable feature dimensions (e.g. insect antennae, legs and pincers), whereas our study used shapes that varied along a continuum where feature dimensions were not easily separable.

While selective attention to category-relevant features is generally proposed for objects with separable feature dimensions, similar mechanisms may be at play within a single feature dimension^18^. Feature-based attention is known to modulate representations in lower-level sensory areas^53–57^. Traditionally, attention was thought to enhance target representations, but more recent work has shown that, depending on the task, attentional gain may be allocated to off-target neurons in a way that changes the response profile of the network for a specific task context^58–62^. By a similar logic, applying gain to neurons tuned toward the category center, rather than to the target itself, would widen the gap between the neural response to a probe from the same relative to the opposite category and boost the ability to detect a category match. In our study, it is possible that attentional gain to neurons tuned to the center of the category in the DMC task produces the category-dependent shift in the neural pattern we observe thereby setting up a task-specific code for efficiently computing the appropriate decision at test.

Another possibility is that representations of the WM target are unchanged, but that they are accompanied by co-existing representations of the category prototype. This would similarly result in greater similarity within than across categories in the neural population response. Indeed, previous studies have shown co-existing representations of exemplars and prototypes during category learning, albeit in different neural areas^63,64^. Similarly, information in WM may be represented at multiple levels of abstraction along the cortical hierarchy^65,66^. It is worth noting, though, that co-existing representations without any warping of the neural stimulus space, would not explain why we saw reduced generalization in the stimulus code across tasks. Moreover, fMRI studies have observed category-dependent changes within early visual cortex^30,31^, suggesting that stimulus representations themselves are reconfigured, at least in some cases.

Interestingly, while our RSA analyses showed stronger category signals in the DMC task, we also observed a lingering influence of category information on neural patterns in the DMS task in the form of significant distance-to-boundary coding and a negative relationship with the category model. The neural data may help explain the surprising effect of category information on behavioural performance in the DMS task, with lower accuracy near the centre of a category and a repulsive bias in match responses. Together these results suggest that participants may have suppressed category information when it was not relevant to the task. Lingering influence of category learning on behaviour when category information is no longer relevant has previously been reported, though typically in the form of an attractive category bias^18,29,36^. The intermixed task blocks in our design might have encouraged a strategy of suppressing category information in the DMS task, resulting in repulsion rather than attraction toward the category. Future studies may help resolve this discrepancy.

In conclusion, we show that neural WM representations shifted from task-general to task-dependent during the memory delay, in line with a dynamic coding framework. Neural activity primarily coded for stimulus identity during encoding but reconfigured over the course of the delay to integrate task and category membership. Finally, we find evidence for category-dependent shifts in the neural representations in the DMC task, where the neural pattern for same-category items is more similar than items across categories. This category-dependent reconfiguration was related to the task-dependency we observed and may help to efficiently produce the appropriate decision-signal at test. Overall, our results show that task demands drive dynamic representational transformations of WM to support upcoming decision-making.

## 3. Materials and Methods

### 3.1 Participants

Forty-two participants took part in the study. Three participants were excluded due to low performance on the WM task (<75% accuracy in one or both tasks). Three participants were excluded due to excessive eye blinks or eye movements during critical periods (>20% of trials). Finally, two participants were excluded due to technical issues with the EEG recording. The remaining 34 participants were included in the analyses (27 female, 7 male). Their age ranged from 18-31 years (M=19.41, SD=2.66). Participants were recruited through the University of Toronto SONA system and received reimbursement of CAD 15 per hour.

The study was approved by the University of Toronto Research Ethics Board (ethics number: 33694) and all participants provided informed consent prior to taking part in the study.

### 3.2 Stimuli & procedure

Participants completed a computerized experiment consisting of a category learning task followed by a WM task. The tasks were programmed in Matlab R2023a and presented in Matlab R2018a using the Psychophysics toolbox version 3^67,68^. The tasks were presented on a 24-inch Asus monitor (1920×1080) and participants were seated 60 cm from the screen.

#### 3.2.1 Category learning task

Participants first completed a category learning task. They were instructed to pretend they were an astronaut arriving on a new planet. They would encounter different animals (shapes) that they were asked to categorise as either ‘Fuboos’ or ‘Juboos’. Twelve shapes from the validated circular shape space (30 degrees apart) were used as stimuli in the category learning task (Figure 1A)^41^. The shape space was divided in half by a category boundary with six shapes in each category. Before starting the learning task, participants viewed the shapes and the category boundary. The category boundary was counterbalanced across participants (six possible).

During learning, a central fixation cross was presented for an average of 1 s (0.8–1.2 ms; jittered across trials) followed by a shape stimulus in the centre of the screen (4.5 degree visual angle [dva]). Participants were asked to report whether the shape was a ‘Fuboo’ or a ‘Juboo’ using the ‘F’ and J’ keys respectively on a QWERTY keyboard. Following a response, the central fixation dot would flash green if the response was correct and red if the response was incorrect (0.2 s). The shape remained on screen for the duration of the response and feedback period.

Participants completed eight learning blocks of 24 trials each. The learning protocol was designed to encourage a strong representation of the category prototype: the four shapes closest to the centre of each category were repeated three times per block, the four shapes one step further from the centre were repeated twice per block and the four shapes nearest to the category boundary were presented once per block. Additionally, shapes were presented sequentially in the first learning block, so shapes nearest to the centre of the categories were presented first and the shapes nearest to the boundary were presented last. In the remainder of learning blocks the shapes were presented in randomised order.

#### 3.2.2 Working memory task

Following completion of the category learning task participants received instructions for the WM task (Figure 1B). In the WM task, 36 shapes from the validated circular shape space (10° apart) were used as stimuli. On each trial, a fixation dot (0.2 dva) would appear on screen for 0.5 s before the WM stimulus (memory target) was presented centrally for 0.15 s. On each trial within a block, the memory target was chosen at random without replacement from the set of 36 shapes, so each of the shapes appeared once per block. The WM stimulus was followed by a mask presented at the same location for 0.05 s. The mask consisted of 180 shapes from the VCS space overlaid. Following the offset of the mask, the maintenance delay lasted 1.5 s followed by a strong visual impulse (‘ping’) consisting of a smooth disc stimulus (9 dva) presented centrally for 0.1 s. ‘Pinging’ has previously been shown to improve the decodability of orientation stimuli using EEG^44,46,69^. The ‘ping’ was followed by another brief delay of 0.4 s before the onset of the probe stimulus. Participants reported whether the probe matched the memory target or not using the ‘C’ and ‘B’ keys on a standard QWERTY keyboard (response mappings counterbalanced across participants). Participants had 2.5 s to respond. The fixation dot would turn green if their response was correct and red if their response was incorrect (or timed out) for 0.2 s. The inter-trial-interval lasted an average of 1 s (0.8-1.2 s jittered across trials).

On different task blocks participants were instructed to compare the probe to the memory target according to one of two task rules: match-to-sample (DMS) or match-to-category (DMC). On DMS blocks participants were instructed to respond ‘match’ if the probe was the exact same shape as the memory target (50% of trials) and ‘non-match’ if the probe was any other shape (50% of trials; selected at random with replacement from the remainder of the stimulus set). On DMC blocks, participants were instructed to respond ‘match’ if the probe belonged to the same category as the memory target (50% of trials) and ‘non-match’ if the probe belonged to the opposite category. Participants were instructed of the task rule and shown the category boundary prior to starting each block.

There were 864 trials in total divided into 24 blocks of 36 trials. Participants were given the opportunity to take a break after each block of trials. The tasks took about 2 hours to complete in total. Participants practiced before starting the WM task.

### 3.3 EEG set-up and preprocessing

EEG was collected while participants performed the tasks using a 32 channel BrainVision system with a sampling rate of 250 Hz. The data were pre-processed in Matlab using the fieldtrip toolbox^67,70^. The raw data were bandpass filtered from 0.1 to 45 Hz using a Butterworth filter and epoched from-0.5 to 3.2 s relative to the onset of the memory stimulus. Artefacts due to eye blinks were marked automatically using the ft_artifact_eog function (default parameters) on the vertical EOG channel. Artefacts due to eye-movements were marked automatically using custom code identifying deviations >30 mV in the difference between the left and right horizontal EOG channels. Other major artefacts due to drift or muscle movements were marked based on variance in the EEG channels using the ft_rejectvisual function with the ‘summary’ method. We visually inspected the marked artefacts using the ft_databrowser function and adjusted any artefacts that were marked incorrectly using the automatic methods. Trials with artefacts during encoding or delay phases (-0.2-2.2 s) due to eye-movements or identified using the summary method were excluded from analyses. We also excluded trials with eye blinks during WM encoding (-0.2-0.2 s from stimulus onset) and ping presentation (1.7-1.8 s).

### 3.4 Analyses

#### 3.4.1 Behaviour

##### 3.4.1.1 Category learning

The variables of interest were categorisation accuracy and response times on correct trials (RT; in seconds). Mean accuracy and median RT were calculated for each participant as a function of task block (1-8) and distance to category boundary (15°, 45° and 75°). We computed separate one-way repeated measures ANOVAs with factors ‘task block’ and ‘distance to boundary’ for accuracy and RT. Where Mauchly’s test of sphericity was violated we report Greenhouse-Geisser corrected p-values (*p*_GG_corr_).

#### 3.4.1.2 Working memory

The variables of interest were WM accuracy and RT on correct trials (in seconds). Mean accuracy and median RT were calculated for each participant as a function of task type (DMS vs DMC) and paired-sample t-tests (two-tailed) were performed to test for task differences. For all t-tests, we report unbiased estimates of Cohen’s d (also known as Hedge’s g) for effect sizes. Additionally, we calculated accuracy and RT as a function of the angular distance between the target and the probe (binned 10-30°, 40-60°, 70-90°, 100-120°, 130-150°, 160-180°, excluding exact match trials) and as a function of the angular distance of the target to the category boundary (binned 5-25°, 35-55°, 65-85°) separately for the two task types. For each task, we ran one-way repeated measures ANOVAs to test for significant effects of target-probe distance and target-boundary distance on accuracy and RT. In a follow-up analysis, to further investigate a significant effect of target-boundary distance on accuracy in the DMS task, we split the DMS trials by the target-probe distance (match: 0, small: <90°; large: >90°).

To further test if DMS judgements were biased by category structure, we checked if false alarms were more likely for non-match probes that were closer to the centre of the category than the memory target (‘toward’) or further from the category centre than the memory target (‘away’). We limited the analysis to small target-probe distance trials where the target and probe were within the same quadrangle.

#### 3.4.2 EEG

##### 3.4.2.1 Decoding stimulus shape

To test whether the patterns of activity in the EEG channels contained information about the shape in memory, we used Mahalanobis distance to calculate the trial-wise distances between the multidimensional activity at each time point for each of the 36 possible shapes, grouped into 12 angle bins distributed around the 360 degree angle space. We performed the analyses on the 27 EEG channels. Pre-stimulus baseline correction was performed by subtracting the mean voltage prior to WM stimulus onset (-0.2 to-0.1 s) from voltages at all other time points. To ensure the target was encoded, only correct trials were included in the decoding analyses.

The decoding procedure followed an 8-fold cross-validation approach to calculate the decoding accuracy of the shape of interest for each trial. The activity pattern of the trials of the testing fold at a particular time-point were compared to the trials of the 7 training folds. The trials of the training folds were averaged into 12 bins relative to the test trial shape, each bin containing trials with shapes within a range of 30° around the circular shape space. As trials were rejected due to errors and artefacts, we ensured equal trial numbers in each bin by random subsampling. We computed the pairwise Mahalanobis distances between the test trials and each of the shape bins using the covariance matrix estimated from trials in the 7 training folds, using a shrinkage estimator^71^. To obtain a visual representation of a tuning curve, the 12 distances were ordered as a function of angular distance, mean-centred, and sign-reversed, so higher values represented greater similarity between the test trial and the training set. This was repeated for all train and test fold combinations. To obtain reliable estimates, the above procedure was repeated 50 times (with random folds each time), separately for 2 sets of bins used for binning the training trials with respect to the test trial (bin centres: 15-345° and 30-360°, each in steps of 30°).

The resulting 100 samples (50 repetitions×2 angular spaces) were averaged for each of the 12 Mahalanobis distance values at each time point.

Finally, to obtain a summary measure of decoding quality, we computed the cosine-weighted means of the tuning curves^44,45^. Higher values reflect greater shape tuning and thereby greater decoding quality for that trial and chance-level is zero. We took the mean over trials, resulting in a single decoding value for each time-point for each participant.

Decoding time-courses were smoothed with a Gaussian kernel (SD=40 ms) for plotting and statistical testing. We tested for significant decoding at each time-point using a cluster-based permutation test (10,000 permutations) to correct for multiple comparisons over time from stimulus onset to probe onset (0-2.2 s). We used a cluster-forming and cluster-significance threshold of *p*<.05.

##### 3.4.2.2 Within and across task decoding of stimulus shapes

To compute within-and across task decoding quality, we repeated the same methods as above with a few modifications. Because the task rule was blocked, we wanted to ensure that any differences in decoding quality within and across task could not be due to closer temporal proximity of trials within a block. For within-task decoding, we therefore made sure that trials within the same block were never included in the training set by using a leave-one-block-out cross-validation procedure. Otherwise, the procedure was the same as described above. We tested for significant decoding within each task and any difference between tasks in an early (0-0.7 s) and late (0.7-1.7 s) time-window of interest using one-sample (one-tailed, >0) and paired samples (two-tailed) t-tests. For completeness we also tested for significant decoding at each time-point using a cluster-based permutation test as above.

Cross-task decoding used independent data for training and testing and consequently we did not use cross-validation for this analysis. We compared within-and across-task decoding quality based on the data used for testing: DMS within (train DMS, test DMS), DMS across (train DMC, test DMS), DMC within (train DMC, test DMC) DMC across (train DMS, test DMC). For improved power, we averaged across the two tasks to compare within-and across-task decoding quality (see *Supplementary Material* Figure S3A for results separately by task). We tested for significant within-task and across-task decoding in the early (0-0.7 s) and late (0.7-1.7 s) time-window of interest using one-sample (one-tailed, >0) and paired samples (one-tailed; within>across) t-tests. For completeness we also tested for significant decoding at each time-point using a cluster-based permutation test as above.

##### 3.4.2.3 Representational similarity analysis

To investigate the representational geometry depending on task demands, we computed RDMs^42^. First, we rotated the shape space to align the category boundary across participants. As above, the 36 shapes were sorted into 12 shape bins for each task condition, creating a total of 24 conditions for the RDM. Pre-stimulus baseline correction was performed as above. Because we found stimulus decoding quality to be similar for correct and incorrect trials (*Supplementary material*; Figure S2), we included all trials in the RSA to improve power.

To compute RDMs, we first calculated the mean activity for trials belonging to each condition, creating a time-by-channel-by-condition matrix. For each time-point, we then computed the pairwise Mahalanobis distance between each of the conditions using the covariance matrix estimated from residual activity after subtracting the within-condition mean^72^, using a shrinkage estimator^71^.

To test whether the neural data coded for stimulus-level and category-level information at each time-point, we constructed hypothetical models coding for the graded stimulus structure, category structure, and distance to the category boundary (Figure 5A). The graded stimulus model had a graded structure with 0 for identical shapes and 1 for shapes 180 degrees away on the circular shape space, in steps of 1/6. The category model had 0s for shapes within the same category and 1s for shapes belonging to the other category. The distance-to-boundary model had 0 for shapes that shared the same absolute distance to the category boundary, 0.5 for shapes one step closer or further from the boundary and 1s for shapes two steps closer to or further from the boundary. The neural RDM matrices were z-scored before fitting the model. We fit a generalized linear model to each time-point of the RDM, with the graded stimulus model, the category model, and the distance-to-boundary model as predictor variables, including a constant term. The models were fit separately for each task. We extracted the beta coefficients for each predictor model at each time point and these were tested for deviations from zero. Time-courses were smoothed with a Gaussian kernel (SD=40 ms) for plotting and statistical testing. We tested for significant stimulus-, category-and distance-to-boundary coding at each time-point using a cluster-based permutation test as above and in the late time-window of interest (0.7-1.7 s) using one-sample t-tests (one-tailed; >0) and for any task differences using paired samples t-tests (two-tailed).

##### 3.4.2.4 Category-dependent representational shift

Finally, we tested if shapes from within a category were represented more similarly than shapes from different categories, controlling for the physical distance between shapes in the circular shape space. To do so, we repeated the same RDM analysis for each of the individual 36 shapes presented in the experiment, for each of the two tasks separately, creating a total of 72 conditions. For each target shape, we identified shape pairs that were matched for their absolute distance to the target in the circular shape space, but one shape belonged to the same category as the target and the other shape belonged to the opposite category. We subtracted within-category distances from across-category distances to get a measure of category-dependent representational shift. If neural distances are greater across than within category, we would expect this measure to be greater than zero. We tested for category-dependent representational shift within each task at each time-point using a cluster-based permutation test as above and in the late time-window of interest (0.7-1.7 s) using one-sample t-tests (one-tailed; >0) and for differences between tasks using a paired samples t-test (two-tailed).

### 3.5 Data and code availability

Custom code used for data preprocessing and analyses as well as raw and preprocessed data is openly available on the Open Science Framework: https://doi.org/10.17605/OSF.IO/UG5K6.

## 4. Acknowledgements

This research was funded by the University of Toronto Faculty of Arts and Sciences Postdoctoral Fellowship Award to FABP; the Natural Sciences and Engineering Research Council (RGPIN-2017-06866 & RGPIN-2024-05727) and the Connaught New Researcher Award to KF; the Natural Sciences and Engineering Research Council (NSERC) Discovery Grant to MLM (RGPIN-2017-06753 & RGPIN-2024-05884) and the Brain Canada Future Leaders in Canadian Brain Research Grant to MLM. Finally, we would like to thank Mark Stokes and Nicholas Myers for initial discussions that inspired this project and Nicholas Myers for helpful comments on a previous version of this manuscript.

## Author contributions

Conceptualization, F.A.B.P, M.L.M., and K.F..; Methodology, F.A.B.P, M.L.M., and K.F.; Investigation, F.A.B.P and O.B., Resources, M.L.M., and K.F.; Formal Analysis, F.A.B.P, Writing – Original Draft, F.A.B.P Writing – Review & Editing, F.A.B.P, O.B., M.L.M., and K.F..; Funding Acquisition, F.A.B.P, M.L.M., and K.F.; Supervision, M.L.M., and K.F.

