## Supplementary Material for "Task goals dynamically reconfigure neural working memory representations"

**Results**

*Stimulus decoding quality for correct and incorrect trials*

We performed the same decoding procedure as described in the main manuscript and saved the results for each trial. We then sorted the data based on task type (DMS vs DMC) and whether the response at the end of the trial was correct or incorrect (Figure S2). We found no difference in decoding quality between correct and incorrect trials on either task in either the early or late delay period ($p$>.246).

*Stimulus decoding searchlight analysis*

We first calculated the Euclidian distance between channel coordinates in the EEG layout. We sorted distance values to identify the six nearest neighbours for each channel. The decoding procedure followed the same steps as in the main manuscript, but instead of performing the decoding for all channels, we repeated the decoding procedure for each EEG channel and its neighbours separately, to get channel-wise estimates of decoding quality. Mean decoding quality across subjects was plotted for each time window of interest using the ft_topoplotER function from the fieldtrip toolbox (Figure S3)(1). We tested for decoding quality greater than chance (zero) within each channel using paired-samples t-tests (one-tailed) and corrected for multiple comparisons across channels using Bonferroni correction ($p$<.05/27).

In both tasks, we saw a similar pattern where initial stimulus decoding quality during encoding was strongest over posterior cortex and only posterior channels survived correction for multiple comparisons (Figure S2, left panel; DMS: T7, C3, Cz, C4, T8, P7, P3, Pz, P4, P8, PO7, PO3, POz, PO4, PO8, O1, O2; DMC: T7, C3, T8, P7, P3, Pz, P4, P8, PO7, PO3, POz, PO4, PO8, O1, O2; $p$<.05/27). In the early delay, decoding quality was still strongest over posterior channels, but decoding was significant over all channels (Figure S2, middle panel). In the late delay, stimulus decoding quality was more distributed over posterior and left frontal channels (Figure S3, right panel; DMS: AF3, F3, Fz, FC5, T7, Cz, P7, P3, P4, P8, PO7, PO3, PO4, PO8, O1, O2; DMC: AF3, AFz, AF4, F3, Fz, FC5, FC1, T7, C3, Cz, T8, P7, P3, Pz, P4, P8, PO7, PO3, POz, PO4, PO8, O1, O2; $p$<.05), although the only channels that survived correction for multiple comparisons were posterior channels in the DMC task (P7, P3, PO7, POz, PO4, PO8, O2; $p$<.05/27). Interestingly, we found no significant differences between the channels contributing to stimulus decoding between the DMS and DMC task in any of the time-windows (uncorrected or corrected).

*Cost of cross-decoding separately by task*

We computed within-task and across-task decoding quality separately for each task (DMS and DMC), before averaging the results in the main manuscript for improved power. In Figure S4A, we show the results separately for when DMS data was used for testing (left; Within: training DMS, testing DMS; Across: training DMC, testing DMS) and when DMC data was used for testing (right; Within: training DMC, testing DMC; Across: training DMS, testing DMC). For both tasks, there was a numerical trend of greater decoding quality in the late delay (0.7-1.7 s) when training the decoder within than across tasks, but this did not reach significance (DMS: $t_{33}$=1.43, *p*=.081, *d*=.320, 95CI>-.103*10$^{-3}$; DMC: $t_{33}$=1.51, *p*=.071, *d*=.314, 95CI>-.735*10$^{-4}$). There was no difference in the cross-decoding cost between tasks (*p*=.960).

*Cost of cross-decoding searchlight analysis*

We repeated the searchlight analysis described above for within- and across-task decoding and averaged across tasks. Figure S4B shows the distribution of within- and across-task decoding quality across channels in the late delay window (0.7 – 1.7 s) as well as the cost of cross-decoding (within–across; right). Within-task decoding quality was distributed across a network of posterior and left anterior channels (AF3, F3, Fz, FC1, T7, C3, Cz, P7, P3, P4, P8, PO7, PO3, POz, PO4, PO8, O1, O2; *p*<.05/27). Across-task decoding quality was specific to a set of posterior channels (P7, Pz, P4, POz, PO4, PO8, O2; *p*<.05), but none of these survived correction for multiple comparisons. Within-task decoding was stronger than across-task decoding in a group of anterior and two left posterior channels (AF3, AFz, AF4, F3, Fz, FC5, FC1, FC2, FC6, T7, Cz, PO7, O1; *p*<.05), but only one of these survived correction for multiple comparisons (Fz; *p*<.05/27).

### Figures

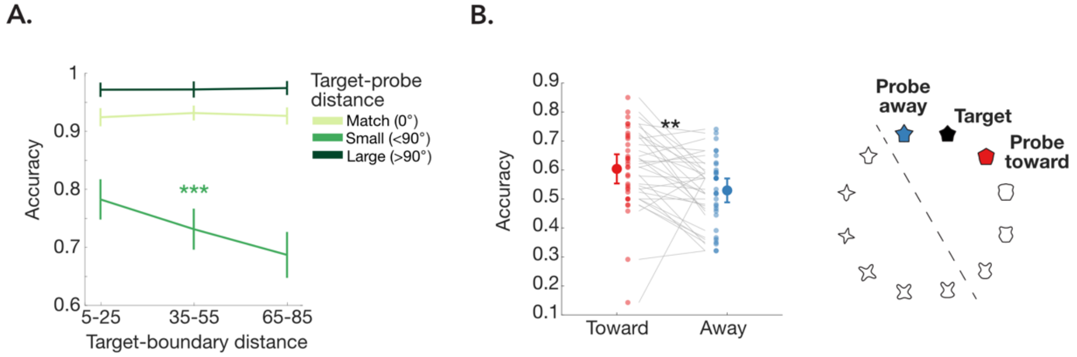


**Figure S1.** Behaviour on DMS task. A. DMS accuracy as a function of the angular distance between the target and the category boundary, split by the angular distance between the target and probe: match trials (light green), small distance <90 deg (medium green), large distance >90 deg (dark green). B. DMS accuracy when target-probe distance was small (<90 deg) separately for when the probe was toward (red) or away (blue) from the centre of the category. Higher accuracy on toward than away trials suggest people are more likely to make an error (i.e falsely endorse a probe as a match) when it is away than toward the category (category repulsion). ***p*<.01, ****p*<.001.

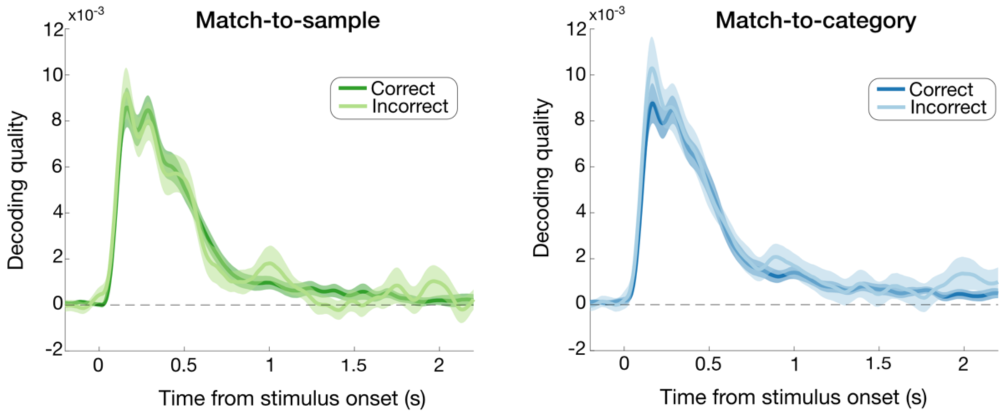


**Fig. S2.** Stimulus decoding quality for DMS (left; green) and DMC (right; blue) task split by correct (dark) and incorrect (light) trials. Shaded area shows standard error of mean (SEM).

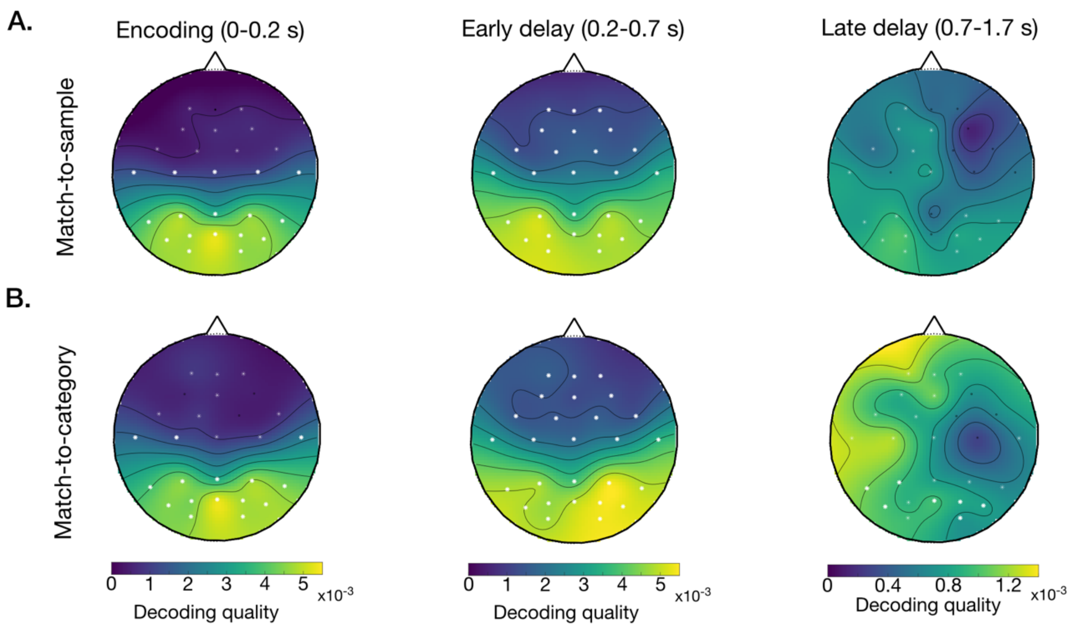


**Fig. S3.** Decoding quality searchlight analysis for DMS task (A) and DMC task (B) averaged across the encoding period (0-0.2s; left), the early delay (0.2-0.7s; midlle) and late delay (0.7-1.7s; right). White stars denote significant channels (*p_uncorrected*<.05). Bolded white stars denote significant channels after correction for multiple comparisons (*p_corrected*<.05/27).

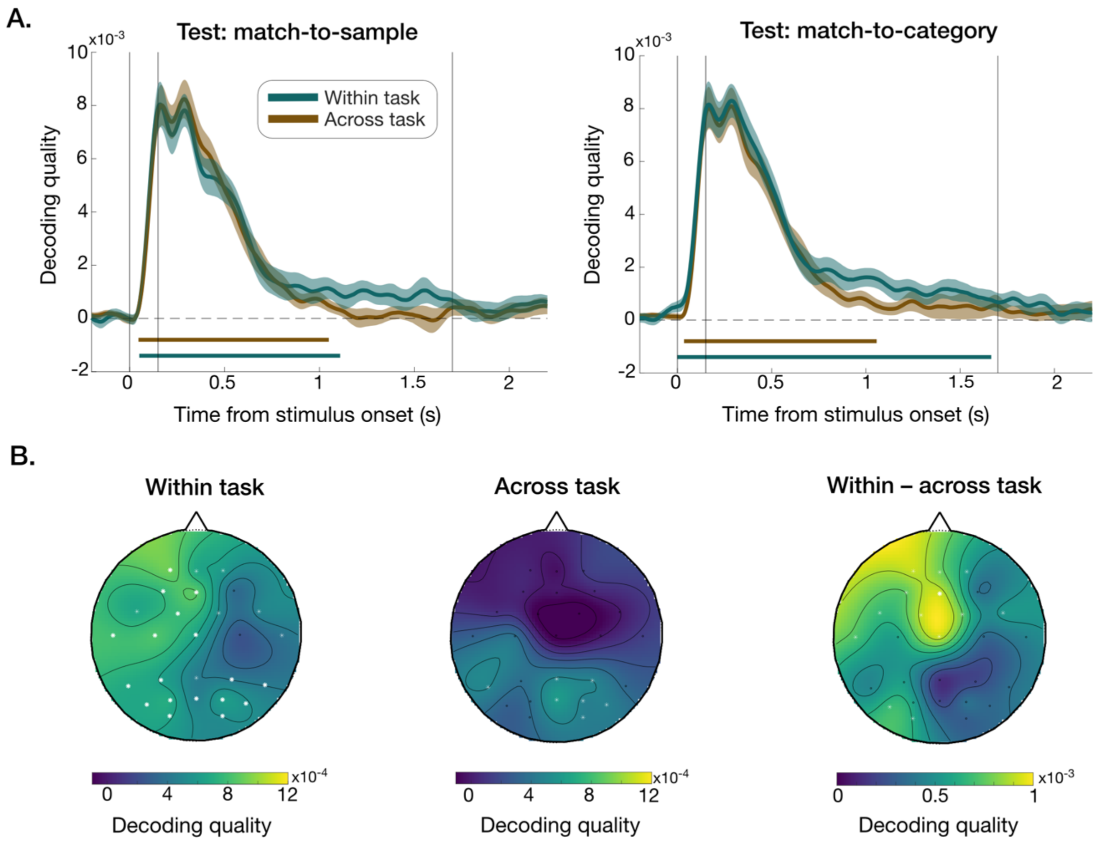


**Fig. S4.** Cost of cross-decoding. **A.** Within- task and across-task decoding quality separately for DMS task (left) and DMC task (right). Shaded areas show SEM. Coloured bars show significant time-points after cluster correction (*p*<.001). **B.** Decoding quality searchlight results of within-task decoding (left), across-task decoding (middle) and within–across task decoding ('cross-decoding cost'; right) averaged across the two tasks for the late delay period (0.7-1.7s). White stars denote significant channels (*p_uncorrected*<.05). Bolded white stars denote significant channels after correction for multiple comparisons (***p_corrected*<.05/27).
